# RadarOmics: Intuitive visualisation of multidimensional omics data in ecological, evolutionary, and developmental studies

**DOI:** 10.64898/2026.01.14.699593

**Authors:** Emma Gairin, Vincent Laudet, Marcela Herrera

## Abstract

Interpreting high-dimensional omics datasets requires visualisation tools that reveal coordinated responses across multiple biological processes. Existing approaches such as heatmaps or enrichment plots typically present processes independently and struggle to convey system-level patterns as experimental complexity increases. We developed RadarOmics, an R package that integrates multidimensional omics data using multi-axis radar visualisations. RadarOmics performs dimensional reduction (based on scaling, Principal Component Analyses, or Linear Discriminant Analyses) for predefined biological processes to extract a representative value for each sample and process of interest. These values are displayed in a circular radar layout, enabling rapid identification of global trends, outliers, and trade-offs among biological functions. Using transcriptomic datasets from anemonefish metamorphosis and zebrafish chemical exposure assays, we show that radar-based visualisations can reveal coordinated molecular responses that are less readily captured with traditional visualisation outputs. RadarOmics provides a flexible and intuitive framework for summarising and interpreting biological variation in ecological, evolutionary, and developmental omics studies, offering a compact system-level overview of molecular behaviour across complex experimental designs.

## 1. Introduction

Modern omics technologies enable genome- and transcriptome-wide analyses across complex ecological, evolutionary, and developmental questions. These approaches generate highly multidimensional datasets—often containing large numbers of samples, treatments, and biological processes—and have transformed our ability to quantify organismal responses to experimental conditions and environmental change. While statistical frameworks for detecting changes in molecular feature abundance and enrichment are well-established, extracting coherent, system-level patterns from high-dimension datasets and presenting them in easily interpretable visualisations remain challenging. Although multiple strategies exist to examine the structure of omics datasets, they each address only part of this issue.

Two main strategies are commonly used to extract patterns in omics datasets. Unsupervised approaches, such as clustering or network-based methods (*e.g.,* Weighted Gene Correlation Network Analysis, WGCNA; Langfelder & Horvath, 2008), identify modules of co-expressed molecular features without prior assumptions, revealing co-regulated networks and novel biological processes. Hypothesis-driven approaches, in contrast, focus on predefined sets of features or functional categories, drawn from *e.g.,* Gene Ontology (GO) terms, KEGG pathways, or curated lists (Herrera et al., 2025). These sets are analysed using enrichment or scoring frameworks, including gene set enrichment analysis (GSEA; Subramanian et al., 2005), gene set variation analysis (GSVA; Hänzelmann et al., 2013), or over-representation analysis (ORA; Khatri et al., 2012). Heatmaps and dimensional reduction techniques, such as Principal Component Analyses (PCA), can also provide visual overviews of profiles for each set. Hypothesis-driven strategies are particularly effective when well-annotated genomes or transcriptomes are available, as they provide direct functional insight. However, both unsupervised and supervised approaches lose interpretability as experimental complexity increases: unsupervised analyses may yield numerous small modules that fragment patterns and obscure biological insight, while supervised analyses are sensitive to noise from co-occurring variables—common in ecological and developmental systems—that produce divergent responses within sets. This loss of interpretability is further amplified in experiments with many samples and treatments, in which case the volume of analyses and visualisations can become overwhelming.

Effective visualisation is key for translating statistical outputs into biological meaning (Gehlenborg et al., 2010; O’Donoghue et al., 2018). Standard graphical outputs, including heatmaps, network plots, and enrichment bar plots, often become cluttered and sequential as the number of treatments grows. Moreover, they typically present results one category at a time, providing little integration across categories and failing to capture how multiple biological functions vary simultaneously across complex experimental or environmental conditions. Ecological, evolutionary, and developmental omics lack a visualisation method that offers a compact, integrative snapshot of system-level patterns, highlighting both consensus and outliers in a single figure (O’Donoghue et al., 2018).

Radar charts offer the kind of multidimensional synthesis that current omics frameworks lack. When group-level patterns are more informative than absolute values, radar charts excel at displaying multiple independent categories simultaneously (Seide et al., 2021). They have become increasingly popular visualisation tools in biomedical fields (Gupta et al., 2021; Saary, 2008; Stafoggia et al., 2011), following decades of use in business, engineering, and computing (Friendly, 2008; Pongswatd & Smerpitak, 2018; Xiao-Tong & Hong, 2017), enabling the rapid detection of temporal changes within individuals or groups, synthesis of multiple outcome measures, and identification of categories most affected by a given treatment (Saary, 2008). However, radar charts remain rarely used in molecular ecology, even though omics datasets possess the precise characteristics that benefit from this type of integrative visualisation: many variables, overlapping processes, and multidimensional responses.

To address this gap, we introduce RadarOmics, an R package designed to integrate and visualise multidimensional omics data using radar charts. RadarOmics applies dimensional reduction to expression or abundance profiles across pre-defined biological categories (*e.g.,* lists of genes from GO terms, KEGG pathways, or other curated process-specific databases) and extracts a representative value for each sample-category combination. These values are displayed in a circular layout, providing a global snapshot of process-level patterns per sample and treatment. RadarOmics highlights outlier treatments, trade-offs among categories, and consensus responses across large datasets in a single integrative figure. Designed to complement pathway- and network-based approaches, it offers an intuitive system-level overview that preserves signals from individual processes. While initially developed for transcriptomic data, this framework can readily be applied to other omics layers—including metabolomics, proteomics, and genomic variation—where features can be grouped functionally. By combining clarity, flexibility, and interpretability, RadarOmics provides a versatile and accessible visualisation framework for exploring the multidimensional nature of molecular responses in ecological, evolutionary, and developmental studies.

## 2. Radar chart computation

RadarOmics is based on dimensional reduction approaches and can therefore handle quantitative, continuous multivariate datasets, such as gene expression matrices, protein abundance matrices, or metabolite profiles. It requires three sets of information (Figure 1A): (i) a matrix of values with samples as columns and variables as rows (*e.g.,* gene expression matrix), (ii) sample information including at least one type of grouping (*e.g.,* experimental treatment), and (iii) a list of variables and their overarching categories (*e.g.,* genes and their corresponding biological processes or modules).

**Figure 1.**
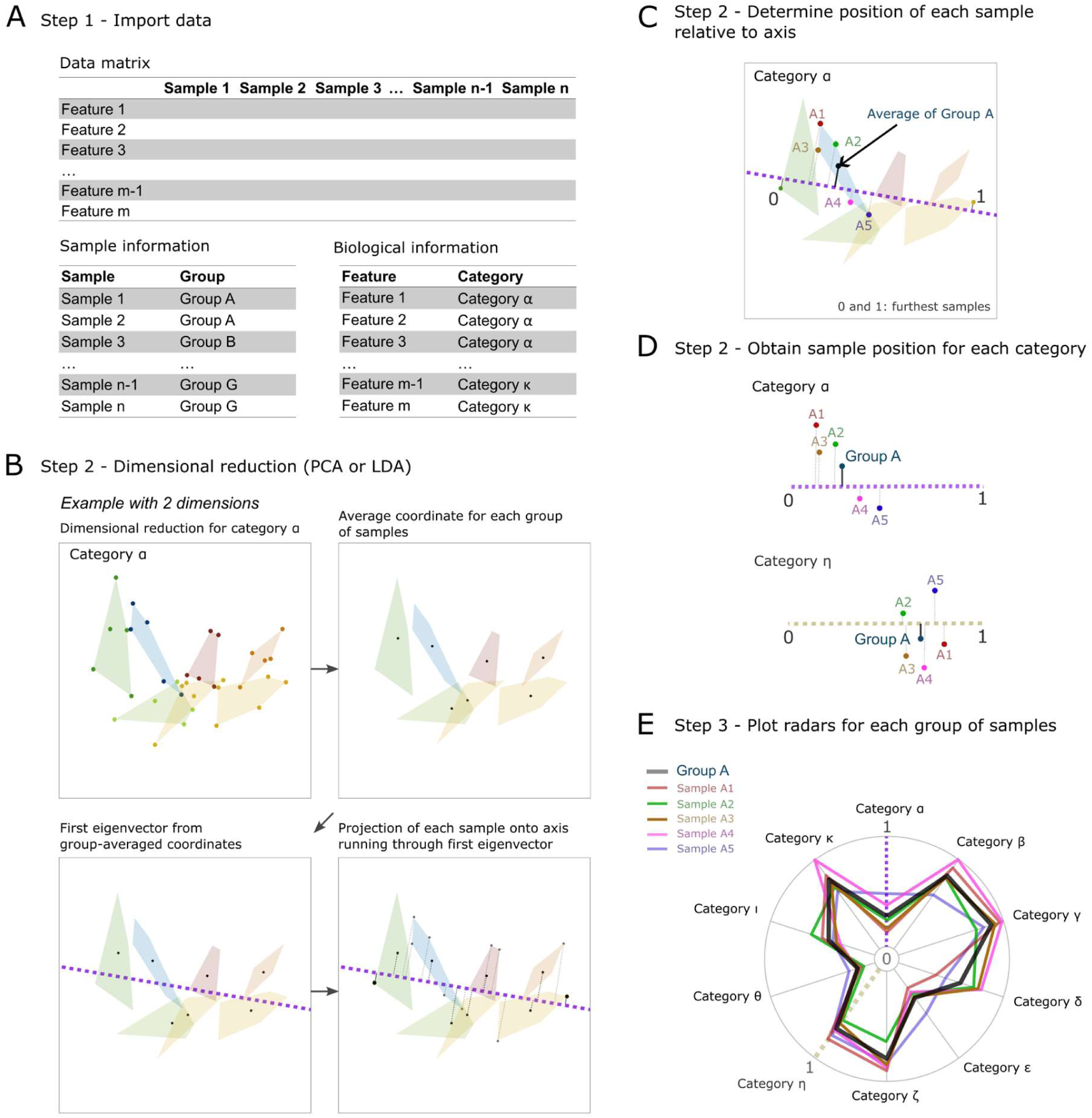
RadarOmics workflow for integrative visualisation of multidimensional omics data using radar charts. A: Step 1: Input data. RadarOmics requires (i) a data matrix containing normalised count or abundance information (*e.g.,* gene expression, protein abundance, metabolite intensity), (ii) sample groupings (“sample information”), and (iii) assignment of molecular features to biological categories (“biological information”). **B:** Step 2: Dimensional reduction. Example using retention of the first two dimensions of a Principal Component Analysis (PCA) or Linear Discriminant Analysis (LDA). For each biological category, coordinates from the retained dimensions are averaged per group, and the first eigenvector of the group-averaged covariance matrix defines the main axis of variance. Each sample is projected onto this axis, producing a 1D representation of its multivariate profile for that category. **C:** Step 2 (continued): Determine position of each sample relative to axis. Zoom-in for category α showing the projection of samples from group A onto the main axis of variance. **D:** Step 2 (continued): Obtain sample position for each category. Extraction of 0–1 scaled coordinates for each category, reducing the multidimensional data to a single representative value per sample. **E:** Step 3: Radar chart generation. Final 0–1 scaled values for each category are plotted in a circular radar chart, where each axis corresponds to a set of features or a biological process (*e.g.,* GO term, KEGG pathway). Coloured lines trace the multivariate position of each sample or group, enabling direct comparison of system-level patterns, outliers, and trade-offs across multiple categories simultaneously.

RadarOmics does not provide options to categorise variables: they must be defined prior to analysis. Lists of features and their biological functions can be derived from manual selection, reference databases such as GO terms, KEGG pathways, modules of co-expression, or curated lists (Herrera et al., 2025). Category size influences the stability of dimensional reduction, so we recommend that biological categories consist of at least 5 biological features.

The data provided to RadarOmics must be transformed so that features are comparable and the variance is stabilised. For transcriptomic datasets, we recommend normalising the data and stabilising its variance before use (*e.g.,* logCPM, calcNormFactors and CPM transformation from edgeR, Robinson et al., 2010; or variance-stabilising transformation and regularised log from DESEq2; Love et al., 2014). For proteomes and metabolomes, datasets can be normalised with median or global scaling approaches and variance can be stabilised with log-transformation.

The main objective of RadarOmics is to extract a single representative value for each sample within each biological category. This value is derived using one of the three dimensional reduction approaches described below. The first option, linear scaling, highlights absolute trends; the second, based on Principal Component Analyses, captures the overall structure without prior assumptions by maximising the variance of the dataset; the third, based on Linear Discriminant Analyses, is supervised and extracts the variation that best separates predefined groups (Figure 2). As LDA assumes that group structure is meaningful, it should be used with care in exploratory analyses.

**Figure 2.**
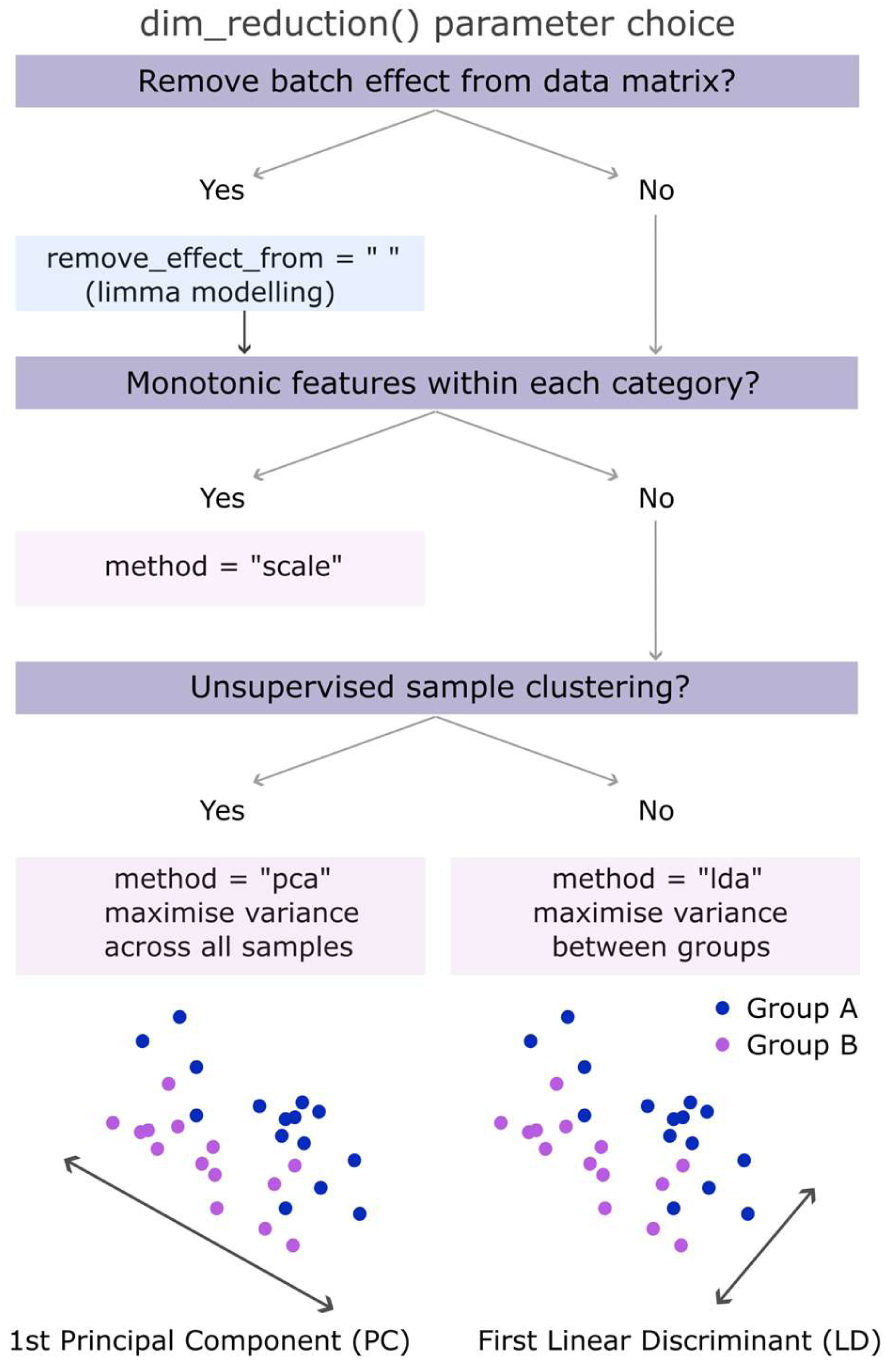
Decision workflow for selecting a dimension-reduction approach and parameters within dim_reduction(). The flowchart outlines the recommended decision process for choosing the appropriate method based on data preprocessing needs and analytical goals. Users first determine whether batch effects should be removed. If biological categories consist of monotonic features (*e.g.,* all genes increasing or decreasing within a given category), scaling is suggested. Otherwise, for unsupervised exploration, PCA identifies axes that maximise overall variance across all samples. When group separation is of primary interest, LDA maximises variance between predefined categories. Example schematics at the bottom illustrate how PCA and LDA identify different optimal axes.

When using linear scaling, for each biological category, the expression or abundance of each molecular feature (*e.g.,* gene, protein, metabolite) is linearly scaled to a value between 0 and 1 (0: lowest expression, 1: highest). The average scaled abundance across all features in a category is then computed for each sample. These averages are linearly scaled from 0 to 1, assigning 0 to the sample with the lowest average and 1 to the highest, which yields a final representative value between 0 and 1 for each sample and category.

The second option is a PCA-based reduction. A PCA is generated using the expression or abundance of molecular features within each pre-defined biological category (Figure 1B). The minimum number of principal components (PCs) required to reach a given portion of the total variance (threshold of *e.g.,* 25, 50, 75%) is assessed. If PC1 meets the variance threshold, the 0–1 scaled PC1 coordinates for each sample are the representative values. If no single PC dominates—a common scenario in ecologically complex of nested designs, multiple PCs are used. To focus on the group-level structure, the average coordinates on these top PCs are computed for predefined groups of samples (note that the choice of how to group samples thus influences the final results). The first eigenvector of the covariance matrix of the group-averaged coordinates is used to define the main axis of variance (Figure 1B). Each sample is then projected onto this axis, yielding points along a segment bounded by the two most extreme samples (Figure 1B,C). These extremes are assigned values of 0 and 1, with the assignment determined from a comparison of the average expression or abundance of molecular features in the two most extreme groups along the axis. Because PCA maximises variance without considering prior groupings, this method faithfully captures the structure of the dataset across categories and groups of samples.

When batch effects distort the PCA structure or more than one PC is necessary to meet the variance threshold, a third option is to use Linear Discriminant Analysis (LDA). Coordinates of each sample across the top PCs are used in an LDA designed to maximise the variance between predefined groups of samples. The minimum number of LDs accounting for a given variance threshold is selected. If LD1 meets the variance threshold, the 0–1 scaled LD1 coordinates for each sample are the representative values. If multiple LDs are required, similarly to the PCA approach, the first eigenvector of the covariance matrix of the group-averaged LD coordinates for the top LDs defines the axis of variance onto which each sample is projected to produce a value between 0 and 1 (Figure 1B,C).

Once a value between 0 and 1 has been defined for each sample and category using any of the three approaches (Figure 1D), RadarOmics produces one radar chart per group, with values near 0 plotted close to the centre and values near 1 at the edge (Figure 1E).

## 3. Implementation in R

Prior to using RadarOmics, three data frames must be prepared in CSV (Comma delimited) format: 1) a multivariate, continuous matrix with samples in columns and molecular features in rows (*e.g.,* gene expression values after variance-stabilising transformation; protein abundances, metabolite intensities); 2) sample information, including sample names and their corresponding group assignments; 3) biological information specifying the molecular feature identifiers (genes, proteins, metabolites, etc.) and their user-defined categories (*e.g.,* manually curated lists, GO terms, KEGG pathways).

The package contains three main functions:

### import_data()

Uploads the data matrix, sample information, and biological information using read.csv() and checks the formatting.

### dim_reduction()

Performs dimensional reduction on the data matrix for each biological category (*i.e.* sequentially using rows corresponding only to genes/proteins/metabolites belonging to each category) and extracts a single representative value per sample and per biological category.

Three dimensional-reduction approaches are available (see Figure 2 for a decision flow chart):

- method = “scale”, recommended for categories with very few molecular features (< 5–10) or when values within the category are consistent or positively correlated.
- method = “pca” (based on prcomp() with scale = FALSE as default, from stats 4.3.3; R Core Team, 2013), recommended for simple experimental designs (*cf.* use example 1) or for exploratory analyses of complex experimental designs where preserving variance across all samples is desirable – particularly when multiple co-variates may have similar magnitudes of effect, or when their relative influence varies across categories.
- method = “lda” (based on lda() from MASS, Venables & Ripley, 2002), recommended when batch effects or strong, non-target covariates influence molecular profiles (*e.g.,* sequencing run, circadian rhythm, developmental stage), or when users want to emphasise group separation in complex designs (see use example 2).

dim_reduction() allows users to specify variance thresholds for PCA and LDA (pca_threshold and lda_threshold). There is also an option to mitigate non-target batch effects via removeBatchEffect from limma (Smyth, 2005), which removes the additive effects of specified nuisance variables (*e.g.,* sequencing run) by regressing them out of the count or abundance matrix. This reduces technical structure while preserving the intended biological signal for dimensional reduction (*cf.* Figure 2).

### plot_radar()

Generates radar charts using the output values from dim_reduction () for each sample and each biological category using ggradar 0.2 (Bion, 2024). One radar chart is produced for each group of samples (default grouping is the “group” column in the sample information file, unless otherwise specified using the argument radar_grouping).

Categories are displayed in the order of appearance from the biological information data frame unless manually specified. Category order can influence how trade-offs, clustering, and directional shifts are perceived on the radar. We therefore recommend grouping related biological functions together (*e.g.,* metabolic processes, signalling pathways) or ordering categories along a biologically meaningful gradient when constructing the plot.

Although plot_radar() supports options for text size and colour coding, fine adjustments to label placement, colours, and background shading are often needed for publication-quality figures. For this reason, we recommend exporting the radar as a PDF or SVG and refining it in vector-graphics software (*e.g.,* Inkscape). This allows users to add visual enhancements such as coloured background shading to group related categories (*e.g.,* energy metabolism, endocrine pathways).

RadarOmics provides several data inspection tools:

- dim_reduction yields an output $information matrix that reports the number of PCs and/or LDs retained for each category and provides the linear correlation (default: Spearman, alternative: Pearson or Kendall) between the extracted values for each sample and the average scaled values of each sample. Spearman correlations are recommended when relationships may be non-linear or influenced by outliers, whereas Pearson correlations are appropriate when linear associations are expected and the representative values for a given category approximate normality. $information thus helps users to evaluate how well the reduced dimensions capture the underlying patterns, which can guide parameter selection and indicate whether manual modifications or closer inspection may be necessary for specific categories.
- plot_dimensions() can be used to visually inspect the dimensionality of the dataset for each biological category by plotting the PCA or LDA space with ggplot2 3.5.1 (Wickham, 2016). It shows how many PCs are required to reach the cumulative-variance threshold and makes explicit when a category is simple (PC1-dominated) or complex (requires several PCs) which helps users to justify parameter choices and understand the structure underlying the radar coordinates.
- plot_boxplot() shows the distribution of the extracted values for each sample and category with ggplot2 3.5.1 (Wickham, 2016) and highlights statistically significant differences between groups of samples using parametric tests (one-way ANOVA and post-hoc Tukey’s HSD tests from stats 4.3.3; R Core Team, 2013) or non-parametric tests (Kruskal-Wallis and post-hoc Dunn’s test with Benjamini-Hochberg correction as default, refer to options provided by dunn.test() from dunn.test; Dinno, 2014) with a threshold p value of α = 0.05 as default. Letters are assigned to each group to indicate significant differences using multcompLetters from multcompView (Graves & Dorai-Raj, 2024). This helps users to quickly identify outliers, bimodal distributions, or categories where heterogeneity may limit interpretability, and ultimately complements the radar plots by communicating to the users that their chosen reduction approach simplifies a potentially complex structure.

Users may combine methods and manually edit the table feeding into plot_radar(). This is useful, for example, when some categories contain too few samples to allow PCA or LDA and must instead be scaled linearly. The pipeline also supports running dim_reduction() on multiple omics layers (*e.g.,* combining RNAseq counts with metabolomics for matched samples), and users may append phenotypic information prior to plotting to obtain radars displaying information from various sources and not only dimensional reduction.

Lastly, RadarOmics is designed to facilitate visualisation across many categories and samples. While flexible, it is intended to complement rather than replace other analytical workflows. For PCA- and LDA-based approaches, we recommend testing multiple variance thresholds and cross-validating category-specific trends with alternative methods (*e.g.,* heatmaps).

## 4. Use examples

To illustrate how RadarOmics complements and extends traditional visualisation approaches such as heatmaps or enrichment plots, we present two representative case studies spanning developmental transitions and toxicant exposures. The first example examines metamorphosis in the false clownfish *Amphiprion ocellaris* (Roux et al., 2023), demonstrating how radar charts succinctly summarise temporal dynamics across multiple pathways and capture stage-specific physiological shifts in a compact, system-level framework. The second example focuses on zebrafish exposed to a multikinase inhibitor (Sorafenib) and a pesticide (Rotenone) across multiple concentrations and developmental time points (Nöth et al., 2025). This case highlights how radar charts integrate multiple treatment-specific responses in complex experimental designs and reveal multidimensional stress signatures that are difficult to synthesise using conventional single-modality plots.

All analyses were conducted in R 4.3.3 and plots were generated using ggplot2 3.5.1 (Wickham, 2016), ggradar 0.2 (Bion, 2024), and ComplexHeatmap 2.18.0 (Gu, 2022; Gu et al., 2016). Statistical significance was defined as p < 0.05 unless otherwise specified. PCAs were performed using the prcomp function and statistical tests included Spearman correlation tests, one-way ANOVA, and linear models (from stats 4.3.3; R Core Team, 2013). Gene counts were normalised with DESeq2 1.36.0 (Love et al., 2014) using variance-stabilising transformation (VST). Manually curated gene sets for *A. ocellaris* and *D. rerio* representing major biological processes – appetite regulation, β-oxidation, brain development, cholesterol synthesis, corticoids, digestion, fatty acid synthesis, gastrointestinal function, glycolysis, ossification, pigmentation, reactive oxygen species, TCA cycle, thyroid cascade, and vision – were used (Herrera et al., 2025). For *D. rerio,* additional categories derived from GO terms were included based on Nöth et al. (2025), such as histone modification (GO terms including “histone”), angiogenesis (“angio”), mitochondria (“mitochondr”), and endoplasmic reticulum processes (“endoplasmic”). All data analysis and figure generation scripts are available on GitHub.

Although the present examples focus on gene expression datasets, RadarOmics is readily applicable to other omics layers—including metabolomics, proteomics, and genomic variation—wherever variables can be meaningfully grouped into functional categories. We emphasise fish datasets as they are the main model taxa in our research group, leveraging our expertise to confidently interpret results and provide detailed biological insights. These systems therefore serve as effective case studies to showcase the versatility and analytical power of radar chart-based visualisations.

### 4.1 Case Study 1: Summarising Temporal Dynamics Across Development

As the first of our two use examples, we applied RadarOmics to a developmental time-series dataset from clownfish (*Amphiprion ocellaris*) comprising 21 transcriptomes across seven post-hatching stages, spanning pre- to post-metamorphosis (Roux et al., 2023; stages defined by Roux et al., 2019). This dataset provides a well-resolved temporal framework, which is ideal for illustrating how RadarOmics condenses coordinated pathway-level dynamics that would typically be dispersed across multiple plots. Raw sequencing reads were processed as in Roux et al. (2023) and variance-stabilised transformation was applied prior to running RadarOmics.

Clownfish development includes a striking metamorphic transition that marks the shift from a pelagic larval lifestyle (stages 1–3), through the thyroid-hormone-triggered onset of metamorphosis (stage 4), and into the benthic juvenile phase (stages 5–7). This transition entails extensive transcriptomic reprogramming, reflected in a PCA based on the 1,000 most variable genes that reveals a clear break at stage 4 and a marked rearrangement thereafter (Figure 3A).

**Figure 3.**
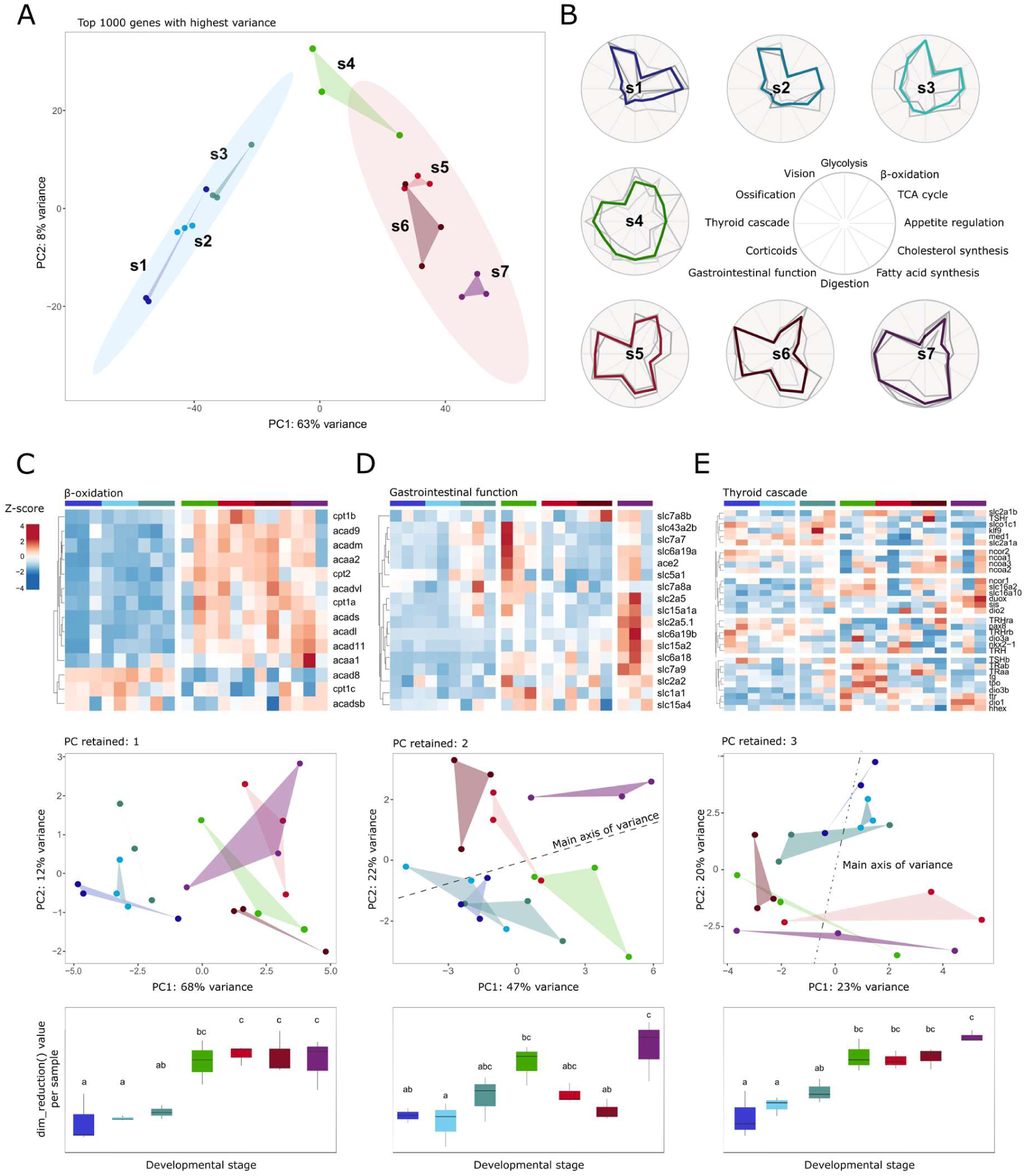
Radar charts capture the complexity of the transcriptomic landscape during clownfish metamorphosis. A: PCA of the 1,000 most variable across developmental stages (s1–s7). Samples are coloured by developmental time, and ellipses grouping stages 1–3 and stages 5–7 were generated using stat_ellipse() in ggplot2 (confidence level = 0.95). B: Radar charts obtained with RadarOmics using the dim_reduction() option set to “pca” (default cumulative-variance threshold = 0.50). Each axis represents a biological category, with values scaled 0–1. C–E: Complementary visualisations produced with RadarOmics functions for each category: C: Heatmap; D: PCA coordinates from plot_dimensions(); E: Boxplot of values from dim_reduction() generated with plot_boxplot(). Categories differ in the number of PCs used to capture variance: C: β-oxidation (1 PC); D: Gastrointestinal function (2 PCs); E: Thyroid cascade (3 PCs; 2 shown).

To illustrate how RadarOmics captures these coordinated shifts, we focused on categories spanning energy metabolism (glycolysis, β-oxidation, TCA cycle, appetite regulation, cholesterol synthesis, and fatty acid synthesis), morphology and physiology (digestion, gastrointestinal function, ossification, vision), and endocrine regulation (thyroid cascade and corticoids). Using dim_reduction() with method = “pca”, pca_threshold = 0.5, and pca_scale = TRUE, we generated one radar chart per developmental stage (Figure 3B), which recapitulated the temporal structure seen in the PCA: gradual shifts stages 1 to 3, an abrupt inflection at metamorphic onset in stage 4, substantial rearrangements through stages 5 and 6, and stabilisation by stage 7. Plotting categories together revealed covariation patterns not readily apparent from univariate summaries. Thyroid-hormone dynamics tracked closely with multiple metabolic and physiological categories (absolute Spearman r = 0.44–0.93, *p* < 0.05 for all except cholesterol biosynthesis), reinforcing its documented role as a system-wide developmental trigger (Roux et al., 2023).

The radar charts illustrated the timing and magnitude of category-specific changes, such as a gradual increase of glycolysis values towards the edge of the plot from stage 1 to 3 before a decline after stage 4 (closely tracking expression profiles, *cf.* Figure 3 from Roux et al., 2023), the steady shift in vision gene expression throughout development, and the three-phase trajectories of digestion, fatty acid synthesis, and corticoid signalling, all of which were stable with low values from stage 1 to 3, increased to medium values between stages 4 and 6, and peaked at stage 7, aligning with gene-level patterns described in Roux et al. (2023).

Beyond corroborating expected trends, RadarOmics highlighted features that may otherwise be overlooked. Ossification values, for example, peaked at stage 6 and diverged from other morphological categories; gene-level inspection confirmed maximal expression of collagen type-I genes (*col1a1a, col1a1b, col1a2*) and periostin paralogues at this stage (Supplementary Figure 1), consistent with a burst of bone-matrix and connective-tissue maturation.

Interpreting some categories requires care: heterogeneous categories may exhibit weak correlations between radar values and underlying expression or abundance patterns. Indeed, radar charts capture multidimensional structure rather than simple average shifts. The TCA cycle, which showed a modest correlation (r = 0.37, *p* = 0.1), illustrated this complexity. Although Roux et al. (2023) reported TCA activation beginning at stage 4, radar values were instead higher and close to edge during early developmental stages here. The $information output of dim_reduction() provides correlations and sign information to guide interpretation and indicates when manual re-orientation may aid clarity.

A final consideration is the number of components retained during dimensionality reduction. Indeed, unlike simply plotting PC1 for each category, the radar values integrate all retained dimensions into a single biologically interpretable coordinate, allowing complex multidimensional patterns to be summarized without discarding informative variance. RadarOmics allows users to specify a cumulative-variance threshold; here we used the default threshold of 0.5, *i.e.,* we retained the minimum number of PCs required to explain at least 50% of the cumulative variance. Categories in which PC1 alone exceeded this threshold used PC1 coordinates, whereas others incorporated multiple PCs. The number of dimensions required reflected the diversity of gene expression within each category (Figure 3C–E). For example, only a single PC was necessary when all genes responded in a tightly coordinated manner, as in β-oxidation (Figure 3C). In contrast, pathways with layered regulation, such as thyroid signalling, required several PCs to meet the variance threshold (Figure 3E). Of the 12 categories examined, 2 were effectively unidimensional (*e.g.,* β-oxidation), 5 required PC1–PC2 (*e.g.,* gastrointestinal function), and 5 required PC1–PC3 (*e.g.,* thyroid signalling). This distribution reflects biological reality: simple, coordinated pathways contrast with cascades that involve multiple regulatory layers. Importantly, even for these higher-dimensional categories, radar profiles captured the expected metamorphic signature from stage 4 onwards, consistent with the thyroid-hormone peak described by Roux et al. (2023) and subsequent shifts in developmental programming.

Together, the combination of dimensionality reduction approaches and radar chart visualisation from RadarOmics efficiently captured and summarised the coordinated transcriptomic reprogramming that underlies clownfish metamorphosis. By condensing complex multi-pathway dynamics into interpretable geometric profiles, the approach highlights how shifts in metabolic and endocrine activity support the ecological transition from pelagic larva to benthic juvenile, offering a clear demonstration of how RadarOmics facilitates integrated, system-level interpretation of temporal omics data.

### 4.2 Case Study 2: Integrating Multidimensional Treatment Effects

While Case Study 1 illustrated RadarOmics’ ability to capture coordinated developmental trajectories, this second case study focuses on a more challenging problem: disentangling overlapping treatment and time effects in a multifactorial design, where ontogeny exerts a dominant influence on gene expression and can obscure treatment-specific signatures. We analysed transcriptomes of *Danio rerio* larvae sampled at 36, 48, 96 hours post fertilisation (hpf) following exposure to Sorafenib (1.3 or 2.4 µM) or Rotenone (25 or 50 µM) from 24 hpf onwards (Nöth et al., 2025). Raw counts from GEO (GSE270785) were transformed into a VST matrix using DESEq2.

We first generated radar charts for each hpf × treatment combination using dim_reduction (method = “pca”) (Figure 4A). Eleven biological categories were chosen based on the results from the original study and to span key metabolic, developmental, and stress-response pathways often disrupted by toxicological exposure, even at early embryonic stages: reactive oxygen species (ROS), mitochondria, endoplasmic reticulum (ER), histone modification, brain development, angiogenesis, ossification, pigmentation, glycolysis, TCA cycle, and cholesterol biosynthesis. Across categories, radar shapes were strongly dominated by time, which was confirmed by linear mixed-effect models showing that hpf explained most of the variance in RadarOmics values, whereas treatment and treatment × hpf contributed little (Figure 4A). Exceptions included mitochondria, brain development, and angiogenesis (Figure 4A,B), consistent with patterns reported by Nöth et al. (2025). Because PCA maximises variance without regard to experimental design, the radar charts in Figure 4A provide an unbiased overview of the dataset’s structure but inevitably mask treatment effects when temporal variation is dominant (*cf.* Supplementary Figure 2). While one could generate separate PCAs for each time point, this would sacrifice the ability to compare developmental trajectories across treatments. Importantly, this highlights a key advantage of RadarOmics over simply inspecting PC1 for each gene set: it offers a structured, category-wise view of multivariate shifts that remain interpretable even when major nuisance effects dominate the dataset.

**Figure 4:**
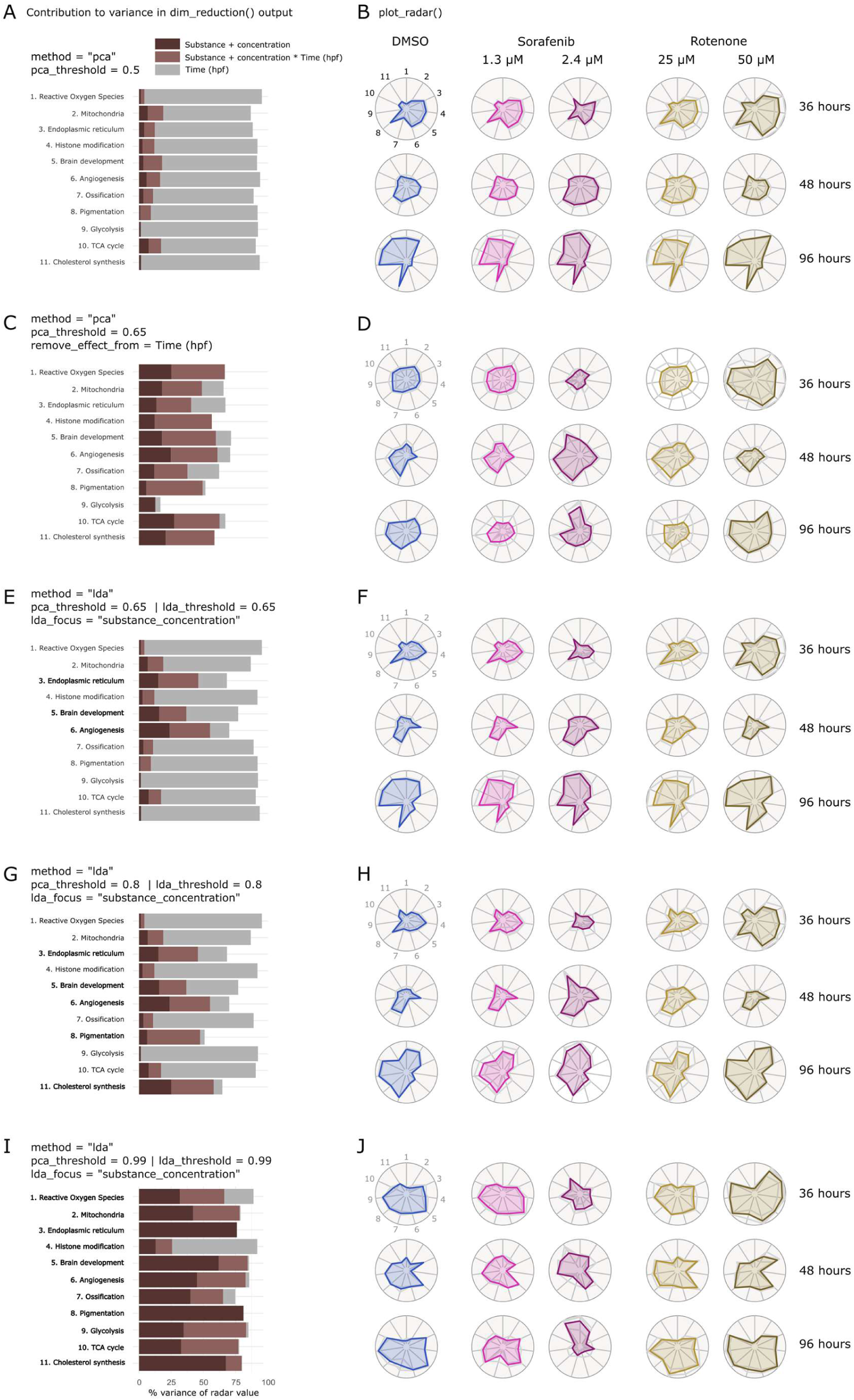
RadarOmics disentangles developmental and treatment effects in a multifactorial zebrafish toxicity experiment. Bar plots (A, C, E, G, I) show, for each category, the percentage of variance in dim_reduction() output attributable to time (hpf), treatment (chemical × concentration), or their interaction under different RadarOmics settings. Under a simple PCA model (A), time explains most variation across categories, whereas treatment and interaction contribute only modestly. Biological processes written in bold underwent LDA, the others were handled with PCA. Corresponding radar charts for each hpf × treatment combination (B, D, F, H, J) reveal a strong ontogenetic footprint, visible as similar shapes within rows across all treatments (B), which can be accounted for through residual fitting (D) or increasing variance thresholds in PCA and LDA (F, H, J).

To address non-target effects such as ontogeny, RadarOmics provides two options: fitting a model and using residuals or performing LDA on category-specific PCs. In Figure 4C-D, we regressed out time (hpf) from the gene expression matrix and generated radars from the residuals using PCA. As expected, the underlying PCA structures (and therefore the resulting radars) differed from Figure 4A-B; however, the rank order of samples within categories was largely preserved (Spearman correlation, *p* < 0.05 for all but cholesterol synthesis). Importantly, removal of the development signal made deviations from control more visually salient within each time step—most notably for the Sorafenib 2.4uM treatment (Figure 4D).

Because residual-based correction may remove meaningful biological signal, we generated three other options of radar plots (Figure 4E-J) to emphasise treatment-induced variation by applying LDA to the set of PC coordinates from each category. As RadarOmics implements LDA on the set of PC coordinates for each category, PCA variance thresholds determine how much information feeds into the LDA step. To test the sensitivity of this pipeline to thresholds and optimise the balance between signal preservation and batch effect correction on a category-dependent basis, we ran LDAs using three PCA/LDA threshold pairs (0.65/0.65 on Figure 4E-F, 0.8/0.8 on Figure 4 G-H, and 0.99/0.99 on Figure 4 I-J), providing progressively more information to the LDA at the cost of diminishing temporal resolution.

For the first pair of thresholds (Figure 4E-F), most categories met the PCA threshold with PC1 alone, meaning LDA was applied to only three categories: angiogenesis, brain development, and endoplasmic reticulum. In these categories, LDAs revealed treatment-specific effects – particularly for Rotenone 50uM at 36 and 48 hpf (Figure 4F) – that were not apparent in the PCA-only radars (Figure 4B, D). At thresholds of 0.8/0.8, five categories did not reach the PCA threshold with a single PC (bold categories on Figure 4G), which markedly increased the portion of variance explained by treatment or treatment and time (Figure 4G). Sorafenib 2.4uM showed striking and consistent deviations from control across categories at all time steps (Figure 4H), while Sorafenib 1.3uM remained similar to control at early time points, reinforcing the temporal and dose-dependent nature of the response. Including nearly all PC information (Figure I-J), on the other hand, minimised the footprint of ontogeny and generated radars that summarised broad differences among treatments. At the most permissive threshold (0.99/0.99), divergences between the two substances became clear: high-dose Sorafenib notably altered ROS, brain development, ossification, glycolysis, and late-stage mitochondria (Figure J) while high-dose Rotenone affected early-stage mitochondrial processes, pigmentation, and the TCA cycle. However, such high thresholds risk overcorrecting or obscuring temporal structure; thus, they should be interpreted cautiously.

This case study demonstrates how RadarOmics flexibly integrates PCA and LDA approaches to reveal treatment-specific molecular fingerprints within nested experimental designs. Exploring multiple thresholds and methods helps to characterise the underlying structure of complex datasets and efficiently highlight biologically meaningful patterns. More broadly, the approach provides an intuitive, category-level summary that can be interpreted not only by domain experts but also by non-specialists, supporting collaboration across multidisciplinary teams.

## 5. Discussion

Modern omics datasets capture thousands of molecular features across complex experimental designs, yet visualising such high-dimensional information in a biologically meaningful way remains challenging. RadarOmics addresses this gap by providing a scalable, flexible, and interpretable framework to summarise multidimensional data across diverse biological contexts. By combining dimensionality reduction with radar-style data visualisation, RadarOmics acts as a magnifying glass to reveal the extent, timing, and impact of treatments on organisms, as well as mechanistic relationships and trade-offs between biological processes. Beyond its novel visualisation output, RadarOmics offers multiple flexible strategies to generate one-dimensional summaries of multivariate omics profiles for each sample. Users can employ simple scaling, unbiased dimensional reduction with PCA, or guided reductions with LDA to maximise variance attributable to target variables and handle strong batch effects. These approaches facilitate compact, interpretable plots that reveal patterns across samples and biological processes, offering a robust alternative to traditional methods that rely on significance thresholds or rank-based scoring.

A key methodological challenge in omics research is extracting a meaningful statistic, score, or value that reflects the profile of a given set of molecular features within a sample or group of samples. Existing approaches—such as significance-based enrichment analyses (*e.g.,* clusterProfiler’s enrichGO) or rank-based statistics (*e.g.,* GSEA or GSVA)—treat all molecular features equally, allowing low-variance features to introduce noise and obscure meaningful patterns. Alternative solutions, including weighted gene scoring (PLAGE; Tomfohr et al., 2005) or transcription factor-based network analysis (VIPER; Cornwell et al., 2018), can provide mechanistic insight but often require extensive pre-processing and may be limited when working with non-model organisms. In addition, although many established tools summarise gene-set behaviour, most rely on statistical significance, p-value ranking, or heatmap-based inspection, all of which can become difficult to interpret in high-dimensional datasets. Heatmaps lose readability as sample numbers increase, PCA biplots saturate with overlapping vectors, and enrichment scores provide group-level summaries but lack per-sample resolution. This gap in interpretability motivated the development of RadarOmics.

By providing a concise representation of process-level activity in each sample, RadarOmics facilitates biological reasoning that is otherwise difficult to extract – such as identifying coordinated or decoupled responses between pathways, revealing early versus late transitions in developmental trajectories, and detecting trade-offs between energetic, metabolic, or regulatory processes. These insights emerge directly from the spatial relationships on the radars rather than from gene-by-gene or molecule-by-molecule inspection. In addition, PCAs and LDAs internally weigh molecular features based on their contribution to variance, downweighing low-variability features, and robustly capturing the overall profile of a given process. A unique feature of RadarOmics is the ability to control the number of retained PCs or LDs based on pre-defined variance thresholds. Unlike approaches that rely solely on PC1 (*e.g.,* ModuleEigengenes in WGCNA, Langfelder & Horvath, 2008; or PLAGE, Tomfohr et al., 2005), RadarOmics flexibly captures multiple-dimensional variance, which is particularly relevant for nested or complex experimental designs.

RadarOmics leverages radar charts, a visualisation format popular in business and diagnostic medicine (Gupta et al., 2021; Saary, 2008) but rarely applied to omics data. By summarising multiple categories in a compact, interpretable format, radar charts allow users to compare large numbers of samples or treatments at a glance. Unlike previous omics applications that plotted individual expression or abundance values along radar axes (*e.g.,* Guthridge et al., 2020; Labaki et al., 2019; Salazar-Noratto et al., 2019), RadarOmics provides robust, biologically meaningful values per sample and category, making the visualisation both informative and versatile. RadarOmics is particularly advantageous for nested or longitudinal designs—such as time × treatment experiments—where traditional visualization approaches struggle to capture coordinated changes across many biological processes. Its ability to summarise each category independently while maintaining comparability across samples makes it well suited for studies with numerous pathways, multi-factor experiments, and analyses in non-model organisms where regulatory networks are incomplete or unavailable. In addition, because the resulting profiles collapse many genes and dimensions into an intuitive visual format, the approach also facilitates communication of transcriptomic patterns to non-specialists, making the dynamics of developmental transitions accessible to broader interdisciplinary fields.

We demonstrated RadarOmics using two transcriptomic datasets, producing visualisations that convey the dynamics, magnitude, and shared responses of molecular features across developmental and treatment conditions. The framework can integrate heterogeneous data types—transcriptomic, proteomic, metabolomic, and phenotypic—on the same visual scale. Users can mix dimensional reduction strategies, include focused molecular feature information (*e.g.,* hub gene expression) or append phenotypic measurements. As with any visualisation framework, RadarOmics has limitations. Radar plots can become visually dense when too many samples are overlaid, strongly correlated categories may produce similar radar shapes, and dimensionality reduction still requires thoughtful selection of variance thresholds. LDA-based reductions also depend on predefined sample grouping. However, these limitations can be mitigated by testing multiple sample groupings, adjusting the number of retained dimensions, or selectively visualising key categories, all of which are supported within the workflow.

RadarOmics is computationally lightweight and scalable, enabling rapid processing of datasets with dozens of categories and hundreds of samples on standard hardware. The framework integrates naturally at multiple points in an analytical pipeline – after QC for exploratory inspection, post–differential expression for summarizing pathway-level responses, or as a final integrative step across multiple omics layers. It is not intended to replace differential expression, clustering, pathway analyses, or rank-based scoring approaches, but rather to complement these methods by providing a coherent visual framework to explore, contextualise, and communicate complex omics results. Researchers can use radar charts to guide targeted follow-up analyses—such as heatmaps or gene-level statistical testing—building an integrated and biologically meaningful interpretation. By extracting system-level patterns while preserving biological meaning, RadarOmics allows researchers to “see the big picture” across complex experimental designs. Its principles extend beyond transcriptomics to multi-omics integration and exploratory analysis, providing a versatile and powerful tool for next-generation biological data interpretation. Future developments may include interactive radar visualisation (e.g., Shiny interface), automated selection of optimal dimensionality reduction strategies, and expanded tools for linking radar outputs back to gene-level drivers. Such extensions will further enhance RadarOmics as an exploratory and integrative platform. To summarise, RadarOmics offers an intuitive, scalable, and flexible framework that bridges the gap between complex multidimensional omics datasets and biologically meaningful, interpretable insights.

## Acknowledgements

Funding for this research was provided by the Japanese Society of the Promotion of Science KAKENHI 24KJ2183. Artificial intelligence tools (ChatGPT) were used solely to support the optimisation of the code used for RadarOmics.

## Data accessibility

The R package RadarOmics is available on GitHub (https://github.com/Emma-Gairin/RadarOmics). All files and codes used in this manuscript are available on Figshare (doi: 10.6084/m9.figshare.31062493).

## Author contributions

Conceptualization: EG, VL, MH

Methodology: EG

Investigation: EG

Visualization: EG

Supervision: VL, MH

Writing – original draft: EG, MH

Writing – review & editing: EG, VL, MH

## Additional information

Supplementary Information is available for this paper. Correspondence and requests for materials should be addressed to Emma Gairin.

---

**Figure S1.**
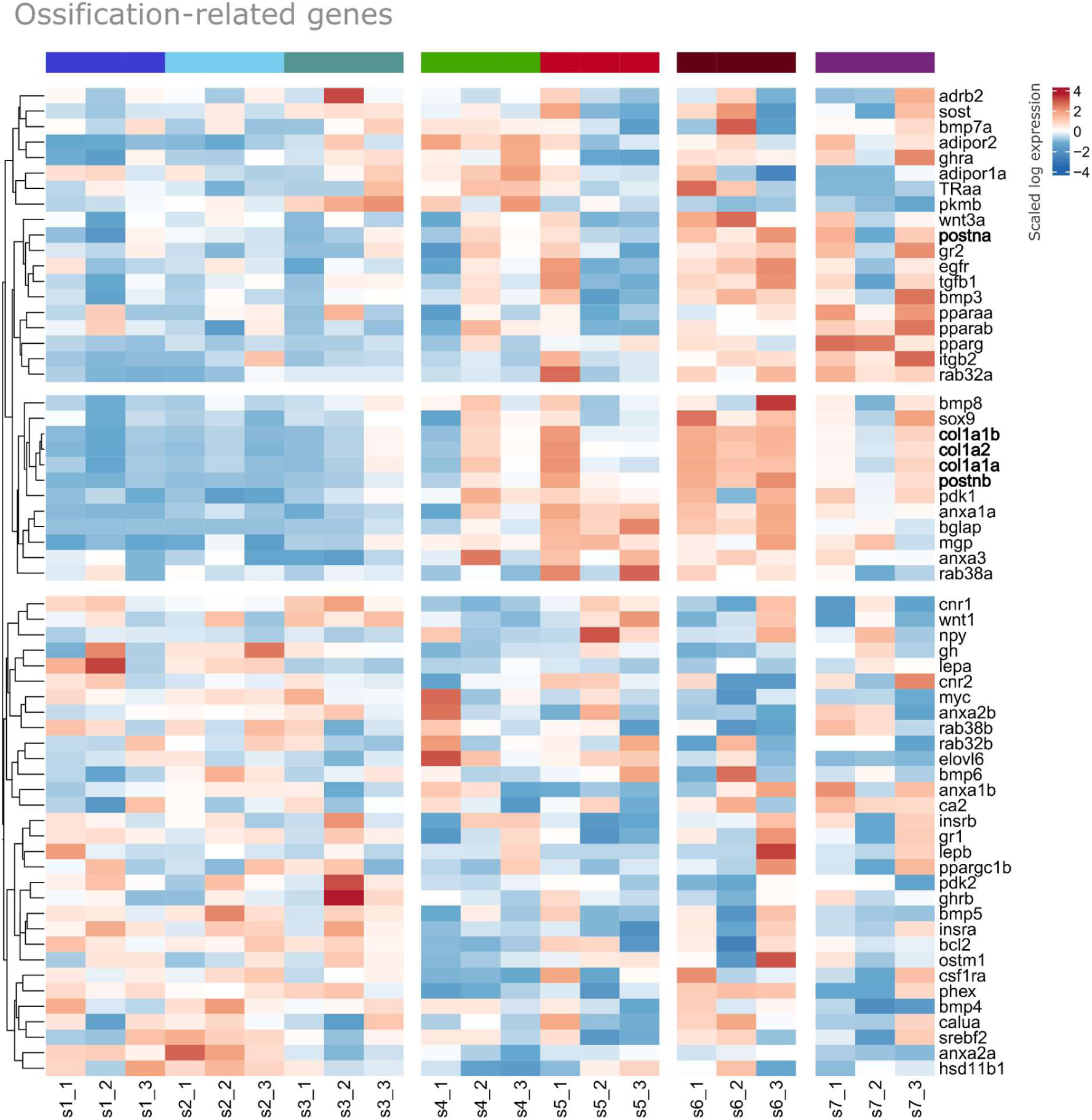
Heatmap of z-score scaled expression level of ossification-related genes (from Herrera et al. 2025) across the seven developmental stages (s1 to s7) of *Amphiprion ocellaris*. Genes highlighted in bold are discussed in the main text.

**Figure S2.**
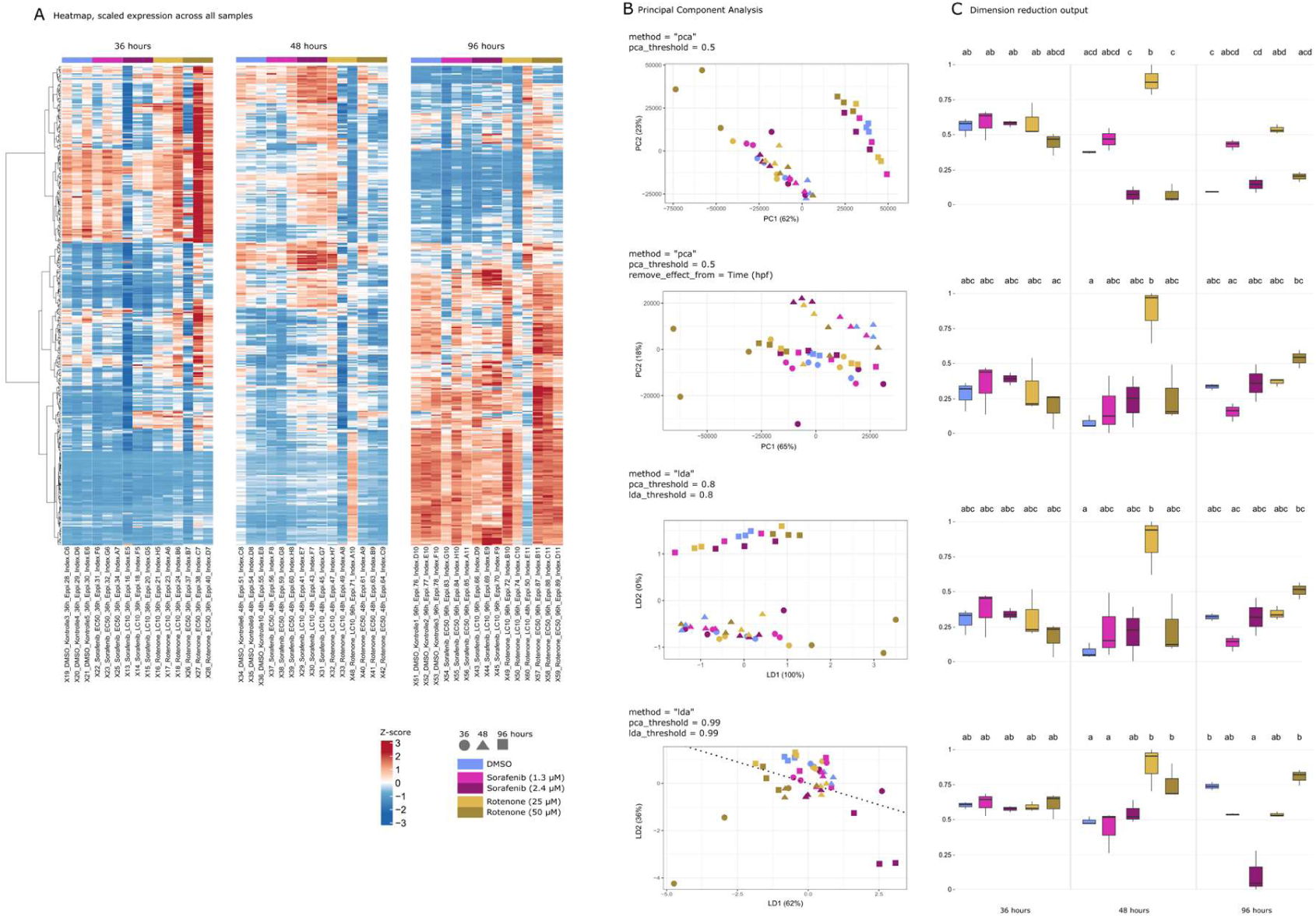
Example of RadarOmics outputs for angiogenesis-related genes in the chemical exposure assay for zebrafish *Danio rerio.* A: Heatmap of z-score scaled expression level of angiogenesis-related genes in the chemical exposure assay for zebrafish *Danio rerio.* B: PCA and LDA produced by RadarOmics (function plot_dimensions()) using the parameters from the main text Figure 4 (method = “pca”, pca_threshold = 0.5 | method = “pca”, pca_threshold = 0.5, remove_effect_from = Time (hpf) | method = “lda”, pca_threshold = 0.8, lda_threshold = 0.8 (same result as for 0.65/0.65 thresholds) | method = “lda”, pca_threshold = 0.99, lda_threshold = 0.99). C: Boxplots produced by RadarOmics (function plot_boxplot()) displaying the representative values extracted from PCA and LDA plots. For all parameters, PC1 or LD1 met the threshold and thus PC1/LD1 coordinates were used, apart for threshold 0.99/0.99 where LD1-LD3 were used. This Figure highlights that Rotenone 25μM stands out at 48 hpf (hours post fertilisation) for any parameter tested. We also observe that remove_effect_from and LDA at moderate thresholds (second and third sets of parameters on panel B) led to very similar structure for angiogenesis. Pushing the thresholds to 0.99/0.99 (fourth set of parameters on panel B) highlighted the absence of significant differences between the treatments at 36 hpf.

## References

Bion, R. (2024). ggradar: Create radar charts using ggplot2. (Version R package version 0.2) [Computer software].

Cornwell, M., Vangala, M., Taing, L., Herbert, Z., Köster, J., Li, B., Sun, H., Li, T., Zhang, J., Qiu, X., Pun, M., Jeselsohn, R., Brown, M., Liu, X. S., & Long, H. W. (2018). VIPER: Visualization Pipeline for RNA-seq, a Snakemake workflow for efficient and complete RNA-seq analysis. BMC Bioinformatics, 19(1), 135. 10.1186/s12859-018-2139-9

Dinno, A. (2014). dunn.test: Dunn’s Test of Multiple Comparisons Using Rank Sums (p. 1.3.6) [Dataset]. 10.32614/CRAN.package.dunn.test

Friendly, M. (2008). The Golden Age of Statistical Graphics. Statistical Science, 23(4), 502–535. 10.1214/08-STS268

Gehlenborg, N., O’Donoghue, S. I., Baliga, N. S., Goesmann, A., Hibbs, M. A., Kitano, H., Kohlbacher, O., Neuweger, H., Schneider, R., Tenenbaum, D., & Gavin, A.-C. (2010). Visualization of omics data for systems biology. Nature Methods, 7(3), S56–S68. 10.1038/nmeth.1436

Graves, S., & Dorai-Raj, H.-P. P. and L. S. with help from S. (2024). multcompView: Visualizations of Paired Comparisons (Version 0.1-10) [Computer software]. https://cran.r-project.org/web/packages/multcompView/index.html

Gu, Z. (2022). Complex heatmap visualization. iMeta, 1(3), e43. 10.1002/imt2.43

Gu, Z., Eils, R., & Schlesner, M. (2016). Complex heatmaps reveal patterns and correlations in multidimensional genomic data. Bioinformatics, 32(18), 2847–2849. 10.1093/bioinformatics/btw313

Gupta, M., Jafari, K., Rajab, A., Wei, C., Mazur, J., Tierens, A., Hyjek, E., Musani, R., & Porwit, A. (2021). Radar plots facilitate differential diagnosis of acute promyelocytic leukemia and NPM1+ acute myeloid leukemia by flow cytometry. Cytometry Part B: Clinical Cytometry, 100(4), 409–420. 10.1002/cyto.b.21979

Guthridge, J. M., Lu, R., Tran, L. T.-H., Arriens, C., Aberle, T., Kamp, S., Munroe, M. E., Dominguez, N., Gross, T., DeJager, W., Macwana, S. R., Bourn, R. L., Apel, S., Thanou, A., Chen, H., Chakravarty, E. F., Merrill, J. T., & James, J. A. (2020). Adults with systemic lupus exhibit distinct molecular phenotypes in a cross-sectional study. eClinicalMedicine, 20. 10.1016/j.eclinm.2020.100291

Hänzelmann, S., Castelo, R., & Guinney, J. (2013). GSVA: Gene set variation analysis for microarray and RNA-Seq data. BMC Bioinformatics, 14(1), 7. 10.1186/1471-2105-14-7

Herrera, M., Vianello, S., Mitchell, L., Chamot, Z., Lorin-Nebel, C., Bonnelye, E., Roux, N., Besseau, L., Gibert, Y., & Laudet, V. (2025). From Genes to Pathways: A Curated Gene Approach to Accurate Pathway Reconstruction in Teleost Fish Transcriptomics. Journal of Experimental Zoology Part B: Molecular and Developmental Evolution, n/a(n/a). 10.1002/jez.b.23299

Khatri, P., Sirota, M., & Butte, A. J. (2012). Ten Years of Pathway Analysis: Current Approaches and Outstanding Challenges. PLOS Computational Biology, 8(2), e1002375. 10.1371/journal.pcbi.1002375

Labaki, W. W., Gu, T., Murray, S., Curtis, J. L., Yeomans, L., Bowler, R. P., Barr, R. G., Comellas, A. P., Hansel, N. N., Cooper, C. B., Barjaktarevic, I., Kanner, R. E., Paine, R., McDonald, M.-L. N., Krishnan, J. A., Peters, S. P., Woodruff, P. G., O’Neal, W. K., Diao, W., … Han, M. K. (2019). Serum amino acid concentrations and clinical outcomes in smokers: SPIROMICS metabolomics study. Scientific Reports, 9(1), 11367. 10.1038/s41598-019-47761-w

Langfelder, P., & Horvath, S. (2008). WGCNA: An R package for weighted correlation network analysis. BMC Bioinformatics, 9(1), 559. 10.1186/1471-2105-9-559

Love, M. I., Huber, W., & Anders, S. (2014). Moderated estimation of fold change and dispersion for RNA-seq data with DESeq2. Genome Biology, 15(12), 550. 10.1186/s13059-014-0550-8

Nöth, J., Michaelis, P., Schüler, L., Scholz, S., Krüger, J., Haake, V., & Busch, W. (2025). Dynamics in zebrafish development define transcriptomic specificity after angiogenesis inhibitor exposure. Archives of Toxicology, 99(4), 1561–1578. 10.1007/s00204-024-03944-7

O’Donoghue, S. I., Baldi, B. F., Clark, S. J., Darling, A. E., Hogan, J. M., Kaur, S., Maier-Hein, L., McCarthy, D. J., Moore, W. J., Stenau, E., Swedlow, J. R., Vuong, J., & Procter, J. B. (2018). Visualization of Biomedical Data. Annual Review of Biomedical Data Science, 1(Volume 1, 2018), 275–304. 10.1146/annurev-biodatasci-080917-013424

Pongswatd, S., & Smerpitak, K. (2018). Applying radar chart for process control behavior. 2018 3rd International Conference on Control and Robotics Engineering (ICCRE), 90–93. 10.1109/ICCRE.2018.8376440

R Core Team. (2024). R: A Language and Environment for Statistical Computing. R Foundation for Statistical Computing. http://www.R-project.org/

Robinson, M. D., McCarthy, D. J., & Smyth, G. K. (2010). edgeR: A Bioconductor package for differential expression analysis of digital gene expression data. Bioinformatics, 26(1), 139–140. 10.1093/bioinformatics/btp616

Roux, N., Miura, S., Dussenne, M., Tara, Y., Lee, S., De Bernard, S., Reynaud, M., Salis, P., Barua, A., Boulahtouf, A., Balaguer, P., Gauthier, K., Lecchini, D., Gibert, Y., Besseau, L., & Laudet, V. (2023). The multi-level regulation of clownfish metamorphosis by thyroid hormones. Cell Reports, 42(7), 112661. 10.1016/j.celrep.2023.112661

Roux, N., Salis, P., Lambert, A., Logeux, V., Soulat, O., Romans, P., Frédérich, B., Lecchini, D., & Laudet, V. (2019). Staging and normal table of postembryonic development of the clownfish (Amphiprion ocellaris). Developmental Dynamics, 248(7), 545–568. 10.1002/dvdy.46

Saary, M. J. (2008). Radar plots: A useful way for presenting multivariate health care data. Journal of Clinical Epidemiology, 61(4), 311–317. 10.1016/j.jclinepi.2007.04.021

Salazar-Noratto, G. E., Nijs, N. D., Stevens, H. Y., Gibson, G., & Guldberg, R. E. (2019). Regional gene expression analysis of multiple tissues in an experimental animal model of post-traumatic osteoarthritis. Osteoarthritis and Cartilage, 27(2), 294–303. 10.1016/j.joca.2018.10.007

Seide, S. E., Jensen, K., & Kieser, M. (2021). Utilizing radar graphs in the visualization of simulation and estimation results in network meta-analysis. Research Synthesis Methods, 12(1), 96–105. 10.1002/jrsm.1412

Smyth, G. K. (2005). limma: Linear Models for Microarray Data. In R. Gentleman, V. J. Carey, W. Huber, R. A. Irizarry, & S. Dudoit (Eds.), Bioinformatics and Computational Biology Solutions Using R and Bioconductor (pp. 397–420). Springer. 10.1007/0-387-29362-0_23

Stafoggia, M., Lallo, A., Fusco, D., Barone, A. P., D’Ovidio, M., Sorge, C., & Perucci, C. A. (2011). Spie charts, target plots, and radar plots for displaying comparative outcomes of health care. Journal of Clinical Epidemiology, 64(7), 770–778. 10.1016/j.jclinepi.2010.10.009

Subramanian, A., Tamayo, P., Mootha, V. K., Mukherjee, S., Ebert, B. L., Gillette, M. A., Paulovich, A., Pomeroy, S. L., Golub, T. R., Lander, E. S., & Mesirov, J. P. (2005). Gene set enrichment analysis: A knowledge-based approach for interpreting genome-wide expression profiles. Proceedings of the National Academy of Sciences, 102(43), 15545–15550. 10.1073/pnas.0506580102

Tomfohr, J., Lu, J., & Kepler, T. B. (2005). Pathway level analysis of gene expression using singular value decomposition. BMC Bioinformatics, 6(1), 225. 10.1186/1471-2105-6-225

Venables, W. N., & Ripley, B. D. (2002). Modern Applied Statistics with S. Springer. 10.1007/978-0-387-21706-2

Wickham, H. (2016). ggplot2: Elegant Graphics for Data Analysis. Springer-Verlag New York. https://ggplot2.tidyverse.org

Xiao-Tong, Y., & Hong, G. (2017). Competitive Evaluation of Internet Enterprises with Improved Radar Chart. 2017 International Conference on Management Science and Engineering (ICMSE), 144–150. 10.1109/ICMSE.2017.8574452

